# Colorimetric CRISPR Biosensor: A Case Study with *Salmonella* Typhi

**DOI:** 10.1101/2024.08.05.606709

**Authors:** Ana Pascual-Garrigos, Beatriz Lozano-Torres, Akashaditya Das, Jennifer C Molloy

## Abstract

There is a critical need to implement a sensitive and specific point-of-care (POC) biosensor that addresses the instrument limitations and manufacturing challenges faced in resource-constrained contexts. In this paper we focus on enteric fever which is a highly contagious and prevalent infection in low- and middle-income countries. Although easily treatable, its ambiguous symptoms paired with a lack of fast, accurate and affordable diagnostics lead to incorrect treatments which exacerbate the disease burden, including increasing antibiotic resistance. In this study, we develop a readout module for CRISPR-Cas12a that produces a colorimetric output that is visible to the naked eye and can act as a cascade signal amplifier in any CRISPR assay based on trans-cleavage. We achieve this by immobilizing an oligo covalently linked to a β-galactosidase (LacZ) enzyme, which is cleaved in the presence of DNA target-activated CRISPR-Cas12a. Upon cleavage, the colorimetric enzyme is released, and the supernatant transferred to an environment containing X-Gal producing an intense blue color. This method is capable of detecting amplified bacterial genomic DNA and has a lower limit of detection (LoD) to standard fluorescent assays while removing the requirement for costly equipment. Furthermore, it remained active after lyophilization, allowing for the possibility of shipment without cold chain, significantly reducing deployment costs.

Scheme 1.
Mechanism of CRISPR-Cas12a, and the signal output produced by enzyme trans-cleavage and the release of LacZ. 1) Initial set-up of inactivated system. 2) DNA isolation and amplification. 3) DNA addition to the biosensor. 4) CRISPR recognition of the DNA target and indiscriminate cleavage of all DNA. 5) LacZ release to the supernatant. 6) Transfer to an X-Gal environment. 7) X-Gal cleavage by LacZ and chromophore oxidation to the intensive blue indigo [illustration created with BioRender.com].

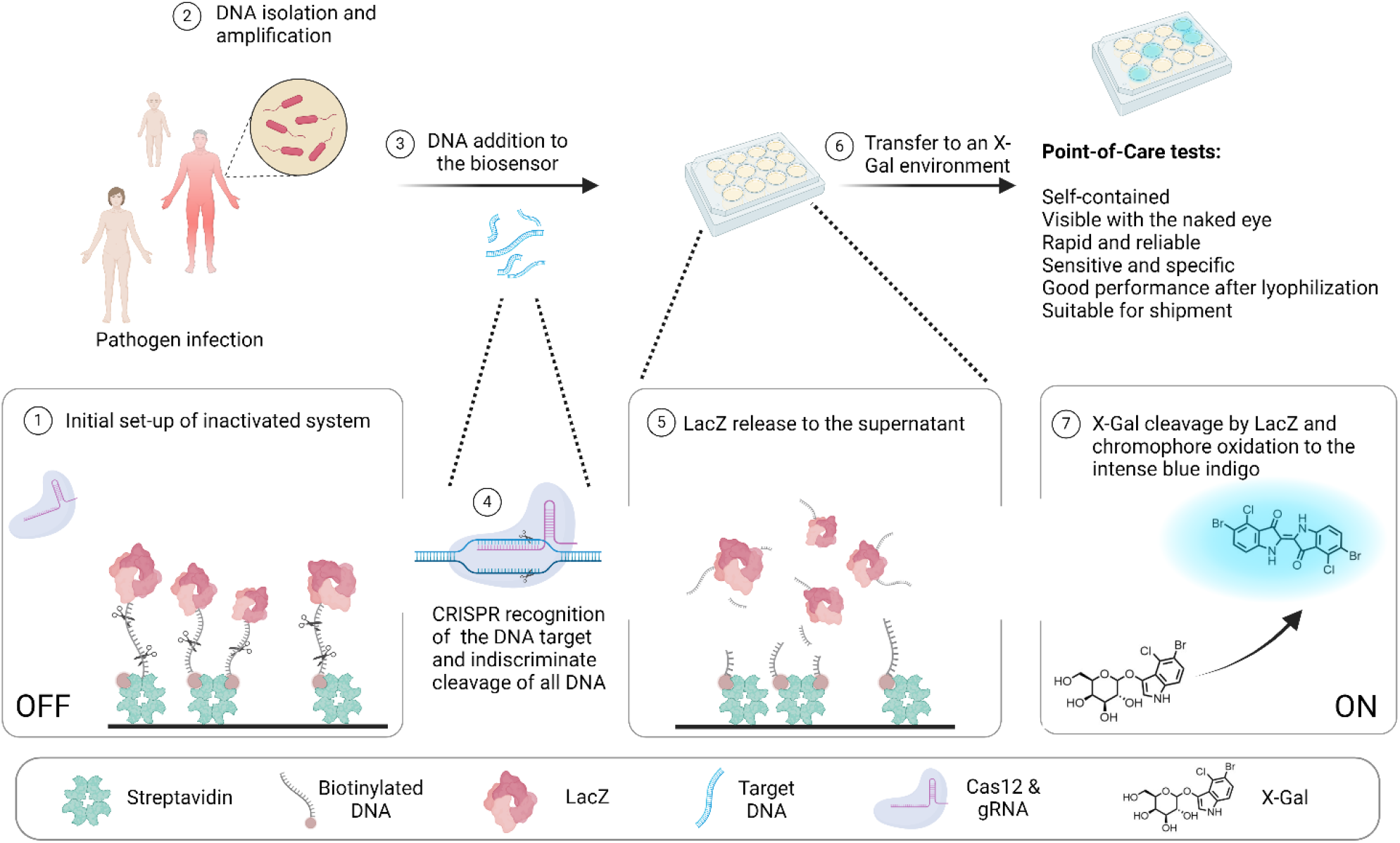

---

Enteric fever is a highly contagious infection caused by *Salmonella enterica* serovar Typhi (*S*. Typhi) and Paratyphi A, B & C (*S*. Paratyphi)^1^. In the last decade, it is estimated that there are 9-17.8 million cases of the disease and around 100,000-208,000 deaths worldwide, but primarily concentrated in resource-limited settings. Infections are usually spread by contaminated food and water supplies^2–5^.

Enteric fever is treated with antibiotics such as ciprofloxacin and ofloxacin^6^. However, it is difficult to diagnose clinically as it presents with non-specific symptoms such as diarrhoea, nausea and abdominal pain.^5,7^ Blood, stool and bone marrow cultures are considered standard practice but take days to confirm a diagnosis^4^. When left untreated, enteric fever can lead to ulceration, bleeding and even multiorgan failure^1,8^. So, the speed of diagnosis is a critical factor in achieving good outcomes following infection.

Concerning the speed of diagnosis of the gold-standard, blood culture, a potential solution is the use of nucleic acid tests (NATs) such as polymerase chain reactions (PCRs) to identify the appropriate *Salmonella* DNA in patient samples. Commonly employed in high-income countries, NATs provide high sensitivity, specificity and quick (1-2 hours) diagnosis. However, these tests are difficult to implement in low- and middle-income countries because of the cost of laboratory facilities and trained personnel^1,9^. A frequently implemented alternative is the Widal test. The Widal test is a low-cost agglutination test that detects the presence of antibodies that are produced in response to enteric fever pathogens. Their sensitivity is generally reported around 80%, while their specificity varies but is consistently reported below 50% due to cross-reacting antibodies from other enterobacteria, among other reasons^10–12^. It is, therefore, not surprising that these values are far from what is accepted by the Foundation for Innovative New Diagnostics’ (FIND) enteric fever target product profiles (TPP)^13^. An ideal test would match the performance of NATs without the need for expensive equipment or trained personnel to perform them as outlined by the REASSURED diagnostics criteria^14^.

A promising approach to an accurate and affordable diagnostic is CRISPR-based NATs. These assays center around a Cas12a enzyme, which is capable of binding to a guide RNA (gRNA) sequence. The gRNA has a hairpin region that binds to the Cas12a enzyme and a spacer region that can be designed to be complementary to the pathogen DNA of interest. Once the Cas12a, gRNA and target DNA form a complex, the enzyme changes conformation to reveal a nuclease domain which indiscriminately cleaves single-stranded DNA (ssDNA)^15^. This indiscriminate cleavage is known as trans-cleavage. By adding in reporters made from ssDNA with a fluorophore on one end and quencher on the other, the indiscriminate cleavage activity results in a fluorescent signal that increases proportionally with Cas12a activity. For low resource settings, one drawback of this

## 2. Materials and Methods

### 2.1 β-galactosidase Expression and Purification

The β-galactosidase expression plasmid (Addgene plasmid # 72935) was transformed into BL21 DE3 cells (NEB). Individual colonies were picked and grown in luria broth (LB) containing 50 μg/mL carbenicillin (Melford) at 37 °C. After 16 hours, the liquid culture was diluted 1:50 into 350 mL of LB-carbenicillin broth and incubated at 37 °C with shaking at 225 rpm until the OD_600_ was between 0.4 and 0.6. At this point, the culture was induced with 1 mM isopropyl β-d-1-reaction is the requirement for specialized equipment to analyze fluorescent readouts^16^.

Colorimetric readouts visible to the naked eye are preferable due to their ease of use and accessibility. Several of these readout methods have been reported in the last five years. A popular method in literature uses CRISPR Cas12a reactions to control the aggregation of gold nanoparticles (AuNPs) by using ssDNA to link the nanoparticles together. A color change from purple to red is observed when Cas12a cleavage allows individual nanoparticles to separate from each other^17–19^. One problem with AuNPs is that the color transitions in many existing assays are subtle and challenging to distinguish reliably with the naked eye, making it difficult for use at POC. Furthermore, AuNPs are prone to aggregation in the presence of biofluids. This can result in false negatives as aggregated nanoparticles will mimic uncleaved ssDNA. They will need to be optimized depending on the nature of the patient sample.^20^

A handful of other approaches have been documented. However, only a minority use colorimetric enzymes and develop a clear color change. A description of these approaches can be found in Table S1. A benefit of using these enzymes is that they act as an amplification step, as each enzyme can sequentially cleave multiple substrate molecules and have potential to improve the limit of detection of the test. These approaches immobilize the enzymes urease^21^ and HRP^22–25^ on a surface with an oligonucleotide. The proteins are released by CRISPR transcleavage activity of activated Cas12a. Then, they are transferred to an environment containing their corresponding chromogenic substrates: urea/phenol red and TMB/H_2_O_2_ producing clear color changes. Inspired by these assays and the scarcity of enzyme-based colorimetric assays, we aimed to develop a system with a clear colorless-to-color reaction, suitable for local manufacturing using a colorimetric enzyme produced in *E. coli*. Additionally, we sought to explore the impact of conjugate DNA length on signal and ensure the system’s compatibility with ambient temperature storage.

In this paper, we propose a detection system using the highly stable LacZ enzyme. Immobilized on a surface with ssDNAs, LacZ is released and mixed with X-Gal, a β-galactosidase-cleavable substrate. The released chromophore spontaneously dimerizes and oxidizes into 5,5’-dibromo-4,4’-dichloro-indigo, producing a colorimetric clear-to-blue output that is visible to the naked eye. We achieve a lower limit of detection (picomolar) than the fluorophore quencher strategy first employed by Chen et al^15^, explore and give insight into the difference made by the oligo linker length^26,27^ and show that our assay can be lyophilized and still maintain functionality. Overall, this tool would improve the diagnosis of enteric fever in the Global South by providing an affordable solution which does not require detection equipment.

thiogalactopyranoside (IPTG) and incubated at 16 °C for another 16 h at 225 rpm. The next day, cells were harvested by centrifugation at 3220 × g for 20 minutes and the pellets were resuspended in 1x PBS and 0.01 mg/mL lysozyme for lysis. The suspended cell mixture was sonicated at high power for 10 cycles of 30 seconds on and 10 seconds off with a MSE Soniprep 150 Plus Ultrasonic Disintegrator (Medical and Scientific Equipment) and clarified by centrifuging at 10,000 x g for 15 minutes. The supernatants were filtered with a 0.2 μm filter and pooled.

The protein was purified through Ni-NTA purification using Ni-IMAC resin (Novagen, #70666-3). The following protocol was adapted from the Novagen His-Bind protocol. After running binding buffer [0.5 M NaCl, 40 mM Tris-HCl, 5 mM imidazole, pH 7.9] through the column and loading the sample, the column was washed with 10-bed volumes of binding buffer followed by 6-bed volumes of wash buffer [0.5 M NaCl, 120 mM imidazole, 40 mM Tris-HCl, pH 7.9]. The protein was finally eluted after applying 6-bed volumes of elution buffer [1 M imidazole, 0.5 M NaCl, 20 mM Tris-HCl, pH 7.9]. The final sample was buffer exchanged into 1x PBS using a CentriPure P25 desalting column (EMP Biotech, CP-0504). The enzyme was concentrated by centrifuging it at 3200 x g for 15 minutes in a Pierce Protein Concentrator PES 10K MWCO (Thermo Fisher Scientific). The expression and purification results were analyzed by SDS-PAGE and glycerol was added to a final concentration of 25 % for storage at -20 °C. Protein concentrations and 260/280 ratios were obtained using a NanoDrop™ One/OneC Microvolume UV-Vis Spectrophotometer (Thermo Scientific) with the parameters: molecular weight (472 kDa) and extinction coefficient (1,046,760).

### 2.2 Conjugation Reaction

DNA oligos with a 5’ amino modifier C12 and a 3’ biotin modification were purchased from IDT containing 20, 40 or 100 nucleotides (nt). The 20 nt oligo was also purchased without the biotin modification (Table S2). All were mixed with 50x excess of Sulfo-SMCC dissolved in DMSO (Thermo Fisher). The final reaction contained 20% DMSO and 80% 10x PBS at pH 8.4 and was incubated at 27 °C with shaking (∼700 rpm) overnight. The sample was then dialyzed for 5 hours in 10x PBS pH 8.4 buffer with a 2 kDa membrane (Fisher Scientific, #15310692) and quantified with a NanoDrop. Oligo-conjugates were characterized by HPLC and mass spectrometry. Next, 25 nmoles of DNA-SMCC were mixed with 5 nmol of purified LacZ so that the DNA was in 5x excess. The reaction was incubated overnight at 27 °C and 700rpm in 3x PBS resulting from the mixing. Dialysis of the sample was carried out in 1x PBS at pH 7 with a 13kDa (SLS, #TUB2002) or 50kDa (Sigma, #PURX50005) membrane, depending on the DNA length, for at least 6 hours.

### 2.3 HPLC

To characterize the oligo-protein reactions, an Agilent 1260 Infinity II was used with a Poroshell 120 EC-C18 2.7 µm 4.6×100 mm (Agilent) stationary phase and a mobile phase containing two buffers: buffer A was made up of 20 mM ammonium acetate at pH 7.7, and buffer B contained 70 % buffer A and 30 % acetonitrile.

For the DNA and DNA-SMCC samples, the protocol ran with 1.3 ml/min flow starting with buffer A at 100 %, then a ramp from 0 to 12 minutes leading to 100 % buffer B and finalized with another ramp from 12 to 15 minutes back to 100% buffer A.

### 2.4 gRNA Design, in vitro Transcription (IVT) and Quantification

During an in-house gRNA screening, spacer regions were selected following a TTTN protospacer adjacent motifs (PAM) sequence. For the target selected for this article, a doublestranded cDNA template was ordered and annealed for gRNA transcription (Table S3). The double stranded products were mixed with IVT reagents from the HiScribe® T7 High Yield RNA Synthesis kit (NEB) and incubated at 37 °C overnight. The gRNA product was purified using DNaseI digestion and a Monarch® RNA Cleanup kit following the manufacturer’s instructions (NEB). The gRNA was further quantified by NanoDrop.

### 2.5 Immobilization of the Conjugates on Streptavidin Plates

384-well Clear Pierce™ Streptavidin Coated High-Capacity plates (Thermo Fisher) were washed 3 times with 100 µl of 1x CutsmartT buffer adapted from the NEB Cutsmart recipe [50 mM Potassium Acetate, 20 mM Tris-acetate, 10 mM Magnesium Acetate, 100 µg/ml BSA, 0.1 % tween-20, pH 7.9]. All conjugates were diluted in the same buffer so that 1 pmol of enzyme monomer was added in 50 µl of buffer unless the concentration was otherwise specified. The conjugate was incubated for 2 hours at room temperature with shaking. After this time, the conjugate solution was removed, and the wells were washed 3 times with 1x CutsmartT. To test successful immobilization, 35 µl of 5 mg/ml X-Gal were added with final buffer contents being 80 % 1x PBS, 10 % DMF and 10 % DMSO to confirm that the conjugates were attached to the plates. The absorbance at 600 nm was collected at 37 °C using a BMG Labtech plate reader. For 96-well streptavidin plates, all volumes were doubled but concentrations remained the same.

### 2.6 CRISPR Trans-cleavage of Immobilized Conjugates

The protocol above was followed for binding up until the second washing step. At this point, 35 µl of CRISPR reaction in 1x CutsmartT were added containing 75 nM LbaCas12a (NEB), 95 nM gRNA and varying concentrations of 60 bp synthetic dsDNA target. Negative controls included reactions without the synthetic target or without the target and gRNA. Benzonase (Sigma) was used as a positive control in the same buffer at 1 unit concentration. The reactions were incubated for 1 hour at 37 °C and the 35 µl of solutions were transferred to a 384-well clear plate (Greiner). 35µl of 5mg/ml X-Gal, as previously described, were added to the cleavage solution. The absorbance at 600 nm was read at 37 °C using a BMG Labtech plate reader.

The CRISPR reaction added in this assay was also tested separately with 1, 0.5 and 0.1 µM of 5’ 6-FAM (Fluorescein)-TTATT-3’ Iowa Black® (FQ) reporter (IDT) and the fluorescence (480 excitation, 520 emission) was measured at 37 °C.

The colorimetric reaction was also tested with PCR amplicons of extracted Typhi DNA (Table S4) and 60 bp synthetic dsDNA off-targets all of which contained a PAM sequence (Table S5).

### 2.7 Limit of Detection Data Analysis

The limit of detection was calculated by normalizing the average value for absorbance at 600 nm of the replicates in each condition at 45 minutes against the average value at 0 minutes. To obtain the % signal, that value was divided by the maximum signal across all conditions and multiplied by 100. These values were plotted against the log10 of the target concentrations used. The slopes of the points before and after signal increase were obtained and the intersection of the two calculated resulting in the limit of detection for each condition.

### 2.8 Typhi Culture, Extraction and Amplification

*Salmonella* Typhi BRD948 (Ty2 ΔaroC ΔaroD ΔhtrA) was grown in 0.2 µm filtered 0.4 mg/ml phenylalanine (Sigma), 0.4 mg/ml tryptophan (Sigma), 0.1 mg/ml para-aminobenzoic acid (Sigma), 0.1 mg/ml dihydro-oxbenzoic acid (Thermo Fisher) and 0.4 mg/ml tyrosine (sodium salt) (Sigma) LB solution overnight shaken at 200 rpm at 37 °C. Genomic DNA was extracted using a Monarch Genomic DNA Purification Kit and quantified using a Nanodrop as above.

### 2.9 Lyophilization

Conjugates were bound and washed as in section 2.5. Then, the CRISPR reaction was mixed as in section 2.6, the only exceptions being that no targets were added, and any Cas dilutions were made into buffers lacking glycerol. The final reactions also contained 10% trehalose dihydrate. The wells were covered in parafilm with a hole above each well and frozen at - 80°C. After one hour, the plate was introduced in a VirTis Advantage lyophilizer and the following protocol was run: -45 °C for 3 hours, -5 °C for 2 hours and 20 °C for 1 hour with a ramp of 0.1 °C/min. The vacuum was held constant at 50 mtorr. After lyophilization, the reactions were rehydrated with and without the target at 7.5 nM and incubated for an hour at 37 °C before the addition of X-Gal as explained in section 2.6.

## 3. Results and Discussion

### 3.1 Conjugate synthesis

Optimization of pH and salt concentration to obtain DNA-NHS conjugates was undertaken using DNA_20_ and fluorescein-NHS (Thermo Fisher). Within the tested range of PBS concentrations (1 x to 10 x), higher concentrations gave higher reaction yields. The effect of volume was also tested but showed no significant difference (Figure S1). Reactions were unsuccessful at pH 7 in all conditions tested but did work at pH 8.4.

Once the reaction was optimized, 20, 40 and 100 nt long aminylated (5’) and biotinylated (3’) ssDNAs were reacted with a cross-linker containing an NHS ester (Figure 1a). The resulting reaction was characterized via HPLC. Compared to unconjugated DNA, the reaction had a higher retention time in the column producing a peak shift for all conjugates (Figure 1b). MALDI-TOF was performed to verify the molecular weight of the reacted species (Figures 1c and S2).

**Figure 1.**
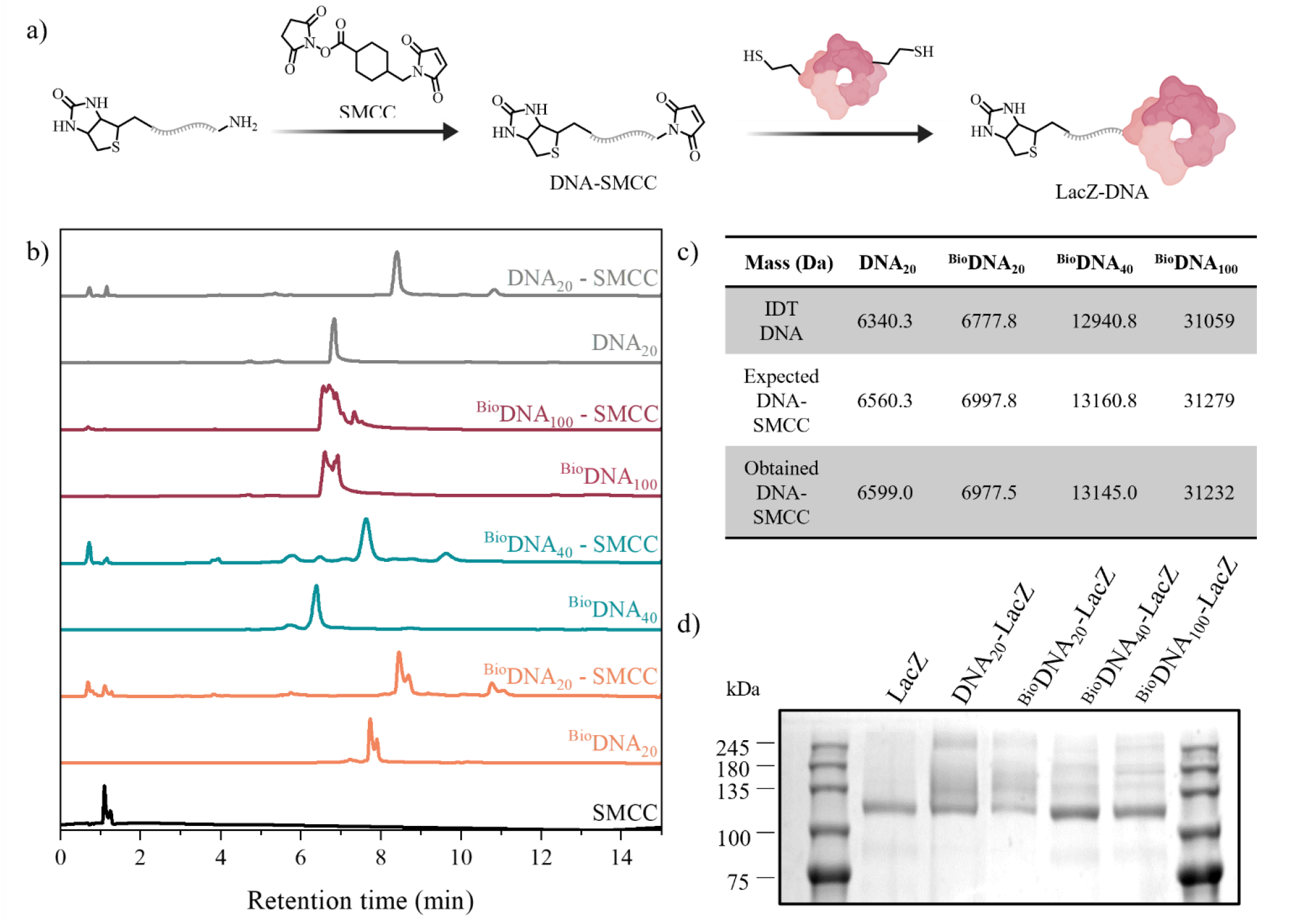
Conjugate characterization. a) Schematic of the conjugation reaction between the aminylated DNA, the SMCC linker and LacZ [illustration created with ChemDraw and BioRender.com]. b) HPLC data of the SMCC linker, aminylated oligos of all lengths and the conjugation of the oligos with the linker. c) Mass spectrometry values of the modified DNA before and after conjugation. d) SDS-PAGE of LacZ at ∼118 kDa vs. DNA-LacZ conjugates at the same and higher masses. The full SDS-PAGE image can be found in figure S3.

Oligonucleotides covalently bound to the cross-linker were incubated in the presence of LacZ to couple the accessible thiols on the enzyme with the reactive maleimide on the linker. After purification, the resulting conjugates were verified by SDS-PAGE which showed bands with higher molecular weights compared to the original LacZ enzyme indicating a covalent modification with the DNA (Figure 1d and S3).

### 3.2 Conjugate binding titration

In order to optimize the conjugate immobilization, the DNA-LacZ conjugates produced were incubated in streptavidin-coated plates at varying concentrations and washed before the addition of X-Gal (Figure 2a). Negative control samples were utilized: unconjugated LacZ and a 20 nt conjugate without biotin (DNA_20_-LacZ). These samples presented negligible binding to the plate whereas the three biotinylated conjugates remained active while bound producing a visible blue color upon addition of X-Gal. The color intensity did not always increase with the concentration of conjugate added, however. We found an optimal concentration between 250 and 1250 fmol of conjugate (Figures 2b and c). It is likely that the higher concentration used inhibits diffusion and reduces binding. Given that the plate manufacturer indicates the binding capacity to be around 60 pmol D-biotin/well, we speculate that the size of LacZ (472 kDa) may be causing steric hindrance, reducing the space available for molecules to bind to streptavidin (60 kDa).

**Figure 2.**
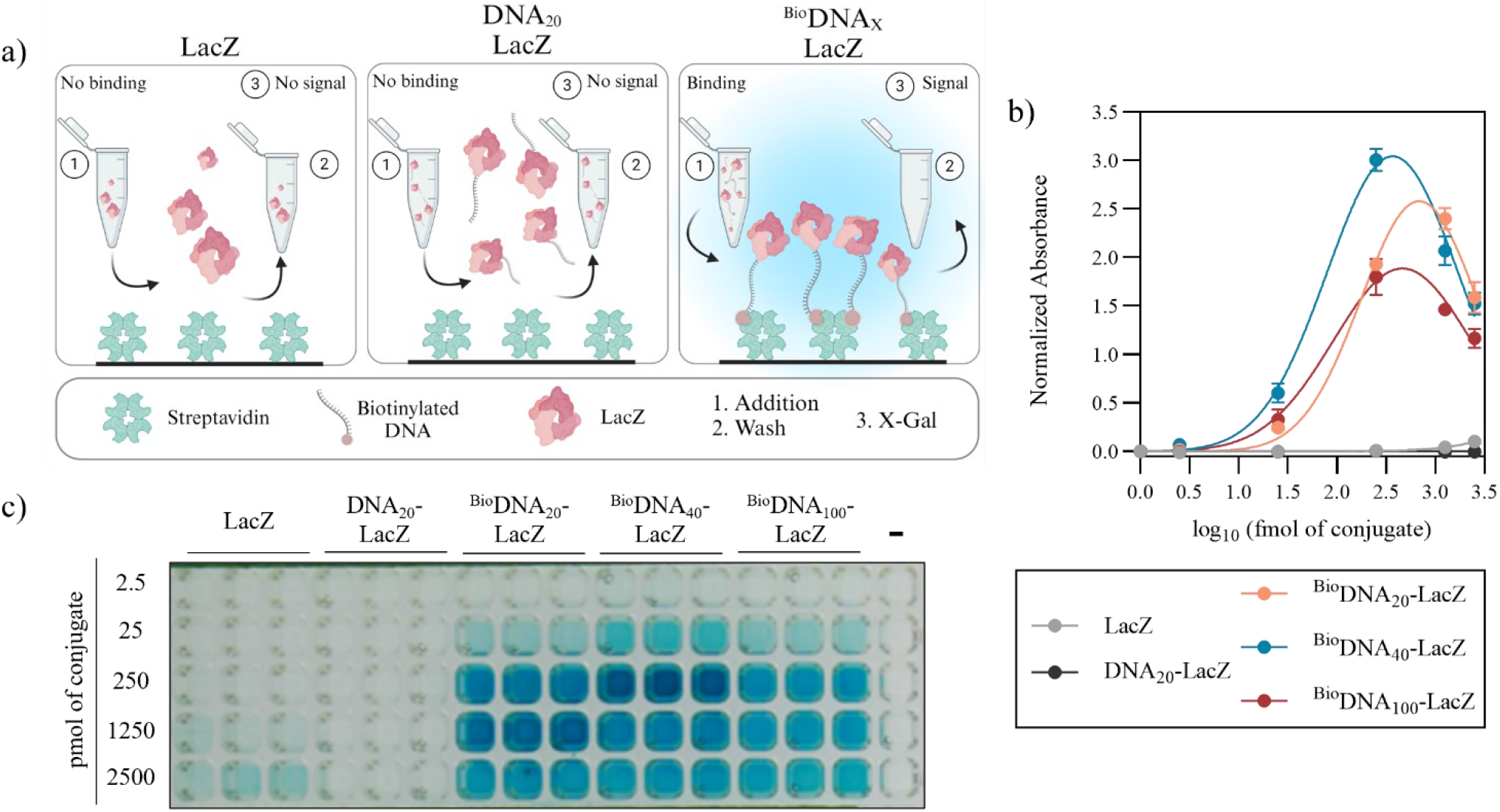
Conjugates Binding Titration. a) Schematic of the two experimental controls which lack biotin, or DNA and biotin, as well as the full conjugates, and the expected colorimetric outcome after binding to streptavidin [illustration created with BioRender.com]. b) Absorbance graph of a titration of conjugates with various lengths of DNA (N=3). c) Photograph of assay results after 45 minutes of incubation (n=3).

### 3.3 CRISPR Conjugate cleavage limit of detection

To colorimetrically detect a *S*. Typhi DNA sequence, we exploited the trans-cleavage property of CRISPR-Cas12a. In the presence of the gRNA/target hybrid, this property was activated and Cas12a cleaved the biotinylated-LacZ hybrid from the streptavidin surface. This produced an intense blue color after transferring the supernatant to a plate containing X-Gal. This was replicated with benzonase in place of Cas12a, acting as a positive control for nuclease activity. In the absence of this target and/or gRNA, the blue color either did not appear or was faint after 45 minutes of incubation (Figure 3a). This time was chosen because background starts to be observed afterwards. Waiting until the reaction plateaued was not feasible due to the hydrolysis product of X-Gal, 5,5’-dibromo-4,4’-dichloro-indigo, precipitating at high concentrations.

**Figure 3.**
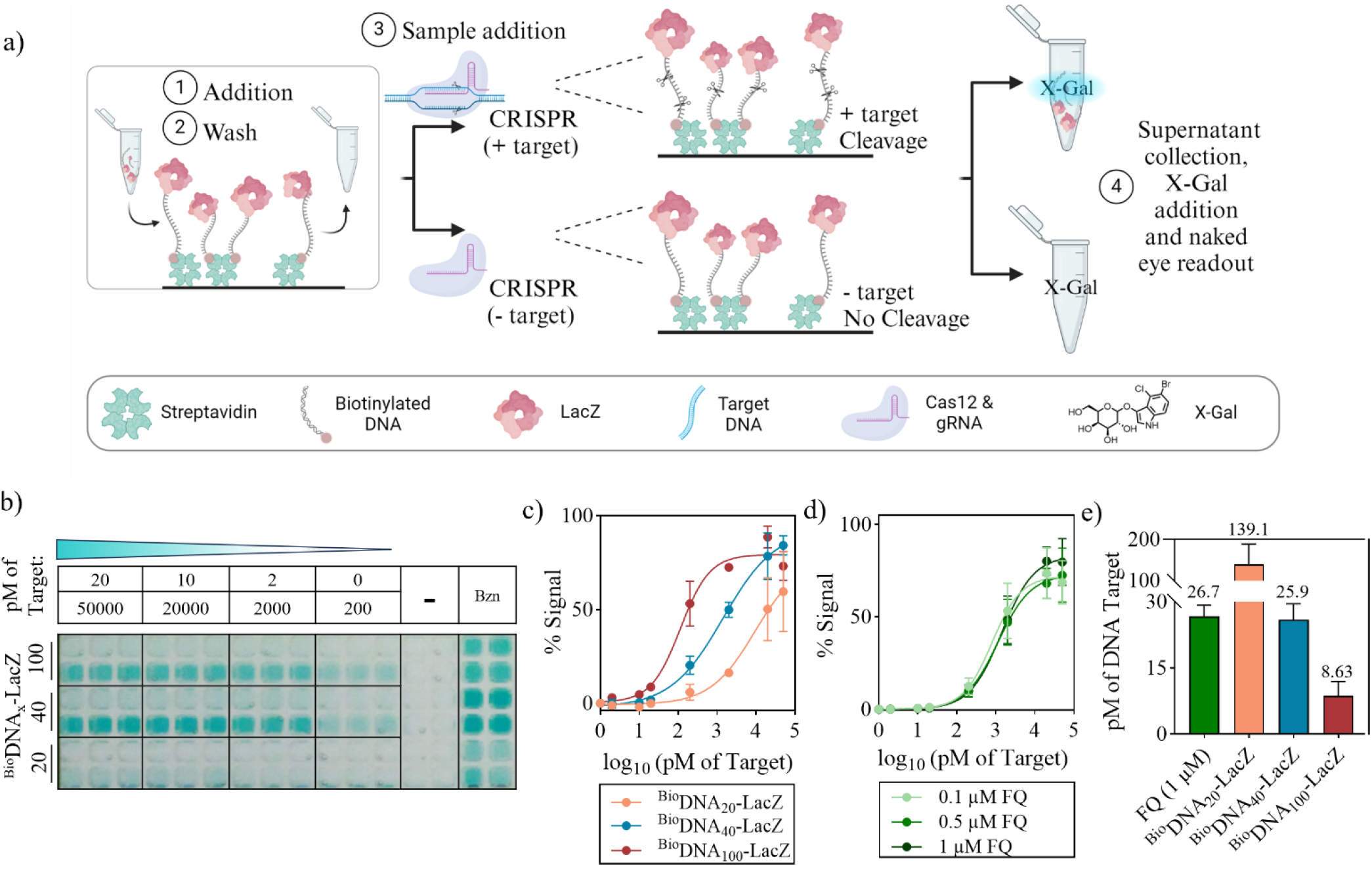
Limit of detection of the CRISPR assay. a) Schematic of the CRISPR assay in the presence of DNA-LacZ conjugates [illustration created with BioRender.com]. b) Photograph of the transferred supernatant following the CRISPR limit of detection assay and incubation with X-Gal for 45 minutes (n=3). c) % Signal from absorbance values of the three conjugates LoD assay in part (b) (N=3). d) % Signal from fluorescence values of the CRISPR limit of detection assay using fluorescent probes at varying concentrations (N=3). e) Calculated limits of detection of each conjugate compared to the fluorescent probe.

A limit of detection study was performed by adding various target concentrations of *S*. Typhi synthetic DNA during the CRISPR reaction step (50 nM, 20 nM, 2nM, 200 pM, 20 pM, 2 pM and 0 pM). After 45 minutes of supernatant incubation with X-Gal, we found limits of detection were inversely proportional to the length of DNA on the conjugate: 139.1 pM, 25.9 pM and 8.63 pM from the shortest to the longest conjugate. The limit of detection was also analyzed visually which increased to 200 pM for the 40 and 100 nt conjugates, and 20 nM for the 20 nt conjugate (Figure 3b and c). From these results it could be hypothesized that an even longer DNA would produce a lower limit of detection by reducing the steric hindrance of Cas12a when allowing more room for the enzyme to reach the DNA between the bottom of the well and LacZ. However, the cost would increase significantly above 100 nt due to DNA synthesis difficulties. Our benchmark CRISPR diagnostic assay used a standard fluorescent reporter that had a LoD of 26.7 pM which is comparable to the 20 and 40 nt conjugates but is significantly higher than the LoD of the 100 nt conjugate of 8.63 pM (Figure 3d and e). P-values for the t-test performed between LoDs can be found in supplementary table S6. Our system thus delivers the benefit of colorimetric readout with higher sensitivity compared to fluorescence, which is typically more sensitive due to lower background noise.

### 3.4 Conjugate cleavage with various DNA sequences

In addition to using synthetic DNA targets, the system was tested with PCR amplified *S*. Typhi genomic DNA. The amplified targets were of various sizes: 500, 2000 and 5000bp (Figure S4). The change in absorbance obtained for all targets was similar with a decreasing trend as DNA length increased but still a clear color difference at 5000 bp compared with the negative control (Figures 4a and 4b). This is expected since the Cas12a would have to parse through more DNA to find its target, slowing it down. This flexibility suggests that this system could be paired with a range of amplification techniques. To investigate sequence specificity of the gRNAs, the system was tested further using short synthetic dsDNA strands that contained identical PAM sequences but with spacer regions that differed from the target. All off-target sequences performed very similarly to the no target reactions as expected (Figures 4c and 4d). Combining all data for target amplicons and off-target samples (Figure 4e), a receiver operating characteristic (ROC) curve shows that the system discriminated well between reactions with and without the correct target sequence with an area under the curve of 1.00 (Figure 4f).

**Figure 4.**
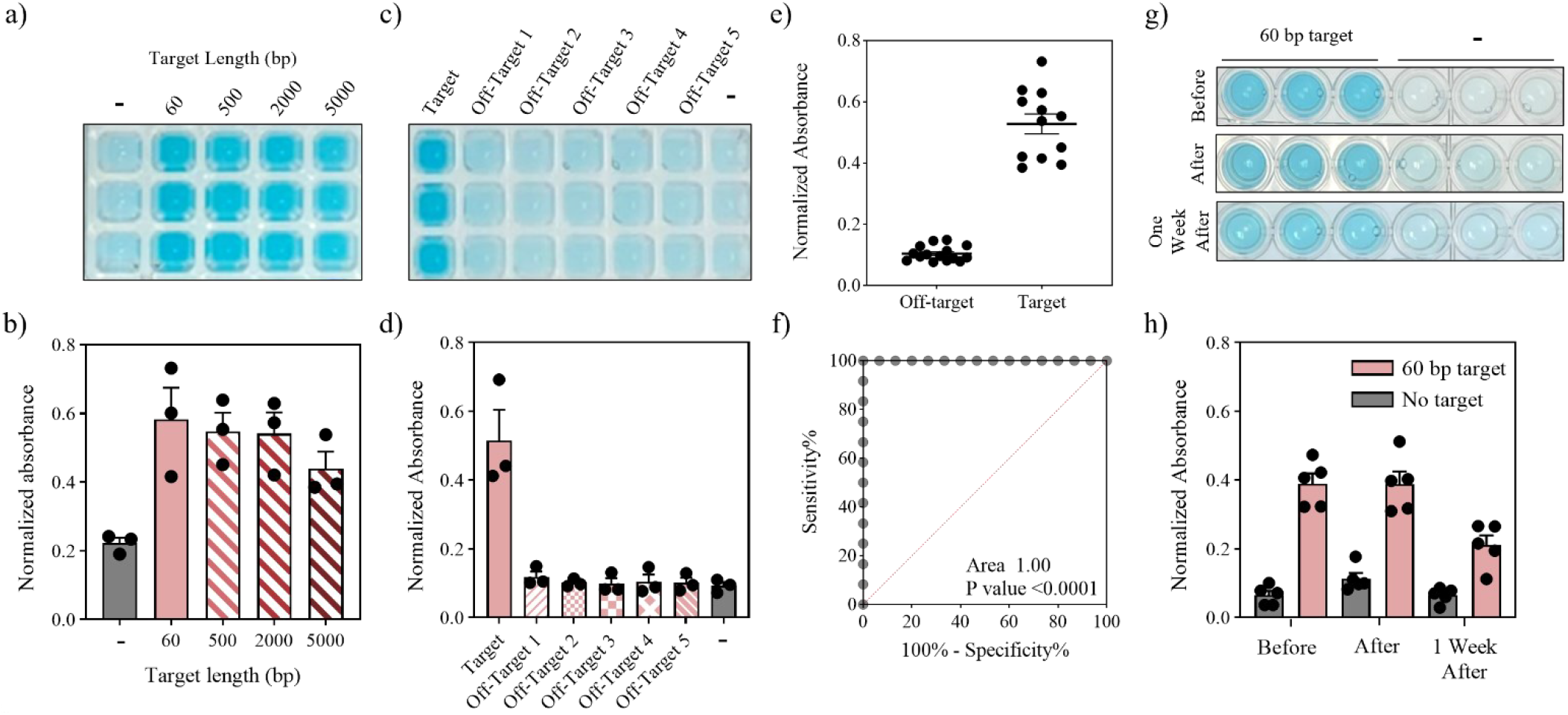
Conjugate cleavage with various targets and lyophilization. a) Image of one replicate of figure (b) after a 45-minute incubation. b) Normalized absorbance after CRISPR cleavage activation with various *S*. Typhi amplicons (N=3). c) Photograph of one replicate of figure (d) after a 45-minute incubation. d) Normalized absorbance after CRISPR cleavage activation with the addition of various off-target sequences (N=3). e) Average values of normalized absorbance after CRISPR cleavage with and without the correct target sequence. f) ROC curve based on the absorbances obtained after 45 minutes of incubation in the presence of target amplicons vs. off-target samples. g) Photograph of part (g) after a 45-minute incubation. h) Normalized absorbance of CRISPR cleavage activation immediately before, after and one week after lyophilization (N=3).

### 3.5 Lyophilization

In most laboratories or clinics, protein-based diagnostics are stored in a fridge or freezer to maintain activity. A cold chain would be necessary during the shipping process which would increase costs. Lyophilization (or freeze drying) would eliminate the need for a cold chain and allow for storage of the diagnostic at room temperature. We tested several lyophilization formulations with the aim of maintaining the structural integrity of LacZ and Cas12a together during lyophilization. Further details can be found in Figure S5.

In our system, the immobilized conjugates were freeze-dried in the presence of Cas12a and a gRNA. After lyophilization, the reactions were rehydrated with and without the target and the absorbance measured. The reactions were also tested before lyophilization in the same conditions to account for the inhibitory effect of trehalose dihydrate. Even in the presence of the sugar, there was a clear color distinction between the samples with target and without. When comparing the activity of the samples before and after lyophilization, there was no significant difference between the two indicating that the enzymes maintained their folded structure and activity. The activity of the system was also tested one week after lyophilization and storage at room temperature. In this case, 57% of the activity remained and a distinct color change was still visible between samples with and without target (Figures 4g and h). The difference was still determined to be significant via a t-test (Table S7).

## 4. Conclusion

This study developed a colorimetric CRISPR system clearly visible to the naked eye with a comparable limit of detection to the gold-standard FQ visualization system. The assay was successful in the presence of *S*. Typhi amplicons, as well as synthetic DNA, with a limit of detection of 8.63 pM, and remained colorless if the target DNA sequence was not present. To extend previous similar, the length of the immobilizing oligos was varied, and we found that the absorbance produced was directly proportional to length. The combined assay with the CRISPR and colorimetric reactions occurred in less than two hours, which is amenable to a POC setting. Lyophilization of the system was successful for one week, so we anticipate that shipping will be possible without cold chain. In future work, optimization would involve coupling the Cas12a detection with an easy amplification technique to reach higher levels of sensitivity, applying the assay to clinical samples, and developing integrated diagnostic devices where the sample preparation, diagnostic assay and readout can be performed in a sequentially. It will also be important to modify this two-step assay to have a one-pot reaction, further simplifying its use. To achieve this, we are exploring enzyme modifications that render the immobilized LacZ enzyme inactive, or that enable inducible activation of the reaction following cleavage. Strategies could include steric hindrance, photocaging, reversible affinity binders or separation of the chromogenic substrate. In addition to detecting *S*. Typhi, our approach would be readily applicable to detection of other nucleic acid targets, and as a readout module for any CRISPR/Cas trans-cleavage based assay.

## AUTHOR INFORMATION

### Present Addresses

Department of Chemical Engineering and Biotechnology, University of Cambridge, Philippa Fawcett Dr, Cambridge CB3 0AS, United Kingdom.

### Author Contributions

APG: conceptualization, methodology, investigation, validation writing – original draft and visualization. BLT: methodology, investigation, writing - review & editing and supervision. AD: conceptualization, writing - review & editing. JM: conceptualization, supervision, writing - review & editing and funding acquisition.

### Notes

The authors declare no competing financial interest.

## ACKNOWLEDGMENT

This research was supported by the mass spectrometry team in the Department of Chemistry at the University of Cambridge. Dr. Derek Pickard (JCBC, Addenbrookes site) provided the *Salmonella* Typhi genomic DNA. We also thank Dr. Steve Baker and Dr. Jim Ajioka for helpful feedback and discussion.

## ABBREVIATIONS

Cas12a: CRISPR-associated protein 12a
CRISPR: clustered regularly interspaced short palindromic repeats
dsDNA: doublestranded DNA
FQ: fluorophore-quencher ssDNA
gRNA: guide RNA
HRP: horseradish peroxidase
HPLC: high-performance liquid chromatography
IVT: in vitro transcription
LacZ: β-galactosidase
LB: luria broth
MALDI-TOF: matrix assisted laser desorption ionization-time of flight mass spectrometry
NAT: nucleic acid test
POC: point-of-care
ROC: receiver operating characteristic
ssDNA: single-stranded DNA
X-Gal: 5-Bromo-4-chloro-3-indolyl β-D-galactopyranoside

