## Supplemental Information for "Colorimetric CRISPR Biosensor: A Case Study with *Salmonella* Typhi"

**Table S1.** Colorimetric approaches coupled with CRISPR-Cas12.

| Method | Model of action | Strengths | Weakness |
|---|---|---|---|
| DNA-conjugated nanoparticle dispersion[1–3] | CRISPR cleavage breaks the DNA sequences responsible for nanoparticle aggregation resulting in a color change as nanoparticles disperse. | One-step reaction | Biofluids can cause aggregation leading to false negatives<br><br>Color change from red to purple may not be obvious<br><br>Linear signal amplification |
| 3,3'-diethylthiadicarbocyanine iodide (DISC2(5))[4] | $DISC_2(5)$ is a colorimetric molecule that is purple in its free form. It can specifically bind consecutive thymidine adenine pairs in dsDNA which causes dimerization of the agent and produces a blue color. In the presence of active Cas12, the dsDNA can be degraded and produce a color change from blue to purple. | No labels needed on the oligonucleotides used, reducing the cost<br><br>One-step reaction | Color change from blue to purple may not be obvious<br><br>Linear signal amplification |
| Platinum nanoparticles[5] | Platinum nanoparticles tethered to magnetic beads by single-stranded DNA are released upon cleavage and travel to a hydrogen peroxide chamber inside a volumetric bar-chart chip. This results in the production of oxygen moving an ink droplet in the device. | Quantitative<br><br>Shows detection of single nucleotide variations | Costly to manufacture the chip<br><br>Linear signal amplification |
| G-Quadruplex DNAzyme[6] | Guanine quadruplexes are complex DNA structures known for their peroxidase-mimicking activity. When intact, they produce color in the presence of ABTS, but if disrupted this activity ceases. | No labels needed on the oligonucleotides used, reducing the cost.<br><br>One-step reaction<br><br>Clear color difference | Counterintuitive color change with green being a negative result and colorless being a positive result.<br><br>Linear signal amplification |
| Fluorophore-quencher probe[7] | Fluorophore-quencher DNA probe as is usually used in fluorescent assays but using a fluorophore which is visible to the naked eye such as ROX. ROX produces a color change from blue to red. | One-step reaction<br><br>Clear color change<br><br>Shown to work with heat from a handwarmer | Photosensitive<br><br>Limitation on how many fluorophores are visible to the naked eye without blue light being necessary |

| | | | |
|---|---|---|---|
| | | | Linear signal amplification |
| Urease-oligonucleotide conjugation[8] | Urease enzymes are tethered to magnetic beads by ssDNA and immobilized. When the enzymes are released by activated Cas12, they are separated and mixed with urea and phenol red. Urea is hydrolyzed and the pH increase produced by the reaction changes the color of phenol red from yellow to red. | Clear color change<br><br>Exponential signal increase | Urea interferant excluding the use of urine samples.<br><br>Biofluids could change the pH, interfering with the assay.<br><br>Two-step reaction |
| HRP-oligonucleotide conjugation [9–12] | An oligo conjugated to HRP and immobilized on a magnetic bead is cleaved by active Cas12. Upon release, the HRP is isolated and mixed with TMB and hydrogen peroxide resulting in the oxidation of the TMB causing an intense blue color. | Clear color change<br><br>Exponential signal increase | Protein difficult to produce in *E.* coli.<br><br>Two-step reaction |
| LacZ-oligonucleotide conjugation (this work) | A DNA-β-galactosidase conjugate was produced and immobilized to a surface. When cleaved in the presence of target-activated CRISPR-Cas12 the colorimetric enzyme is released, and the supernatant transferred to an environment containing X-Gal producing an intense blue color. | Distinct color change.<br><br>Exponential signal increase<br><br>Enzyme easy to produce in *E.* coli.<br><br>Can be lyophilized. | Two-step reaction |

**Table S2.** DNA sequences of the modified oligonucleotides used for conjugation with LacZ.

| | |
|---|---|
| $DNA_{20}$ | /5AmMC12/TCAACGACTGACTGAGCTCT |
| $^{Bio}DNA_{20}$ | /5AmMC12/TCAACGACTGACTGAGCTCT/3Bio/ |
| $^{Bio}DNA_{40}$ | /5AmMC12/TCAACGACTGACTGAGCTCTGATCGAATTCTGACACTACG/3Bio/ |
| $^{Bio}DNA_{100}$ | /5AmMC12/TTTTTTTTTTTTTTTTTTTTTTTTTTTTTTTTTTTTTTTTTTTTTTTTTTTTTTTTTTTTTTTTTTTTTTTTTT TTTTTTTTTTTTTTTTTTTTTTTTTT/3Bio/ |

**Table S3.** Guide RNA cDNA template regions.

| | |
|---|---|
| T7 Promoter | 5'-TAATACGACTCACTATAGG-3' |
| Hairpin recognition region | 5'-TAATTTCTACTCTTGTAGAT-3' |
| Spacer region | 5'-TTAAGAGTGCGTTTGAACAC-3' |

**Table S4.** Primer sequences used to amplify *S.* Typhi DNA samples.

| Amplicon size (bp) | Forward Primer | Reverse Primer |
|---|---|---|
| 500 | CCAGCCCCAACTCAC | ATCATAATCAATATAGCCTTCAGTTTTCCC |
| 2000 | TGGCGCGGGACATGC | TACGTGGTTCTCCGTGTCTGATG |
| 5000 | CGCTGAACTTCCCGC | GACGGACATAATTCAGCGGTAAATC |

**Table S5.** 60 bp Off-Target DNA sequences.

| Off-Target | DNA Sequence |
|---|---|
| 1 | AATATTTAGACTTGCCTTTAAAAGATACCAGAGCCCGAATGACTCGACCATCAGGAACGA |
| 2 | TCGGGCTCTGGTATCTTTTAAAGGCAAGTCTAAATATTGGGAACTTAAGTTTGACCGGG |

| | |
|---|---|
| 3 | ACCCTTGTTAATGCCATTTGGCCCCAGGCTCCCGAGGAGTTATCTGGTAAACTTGCTGCT |
| 4 | TGGACTATGGGACAAGTTTCTTACAGTTGAAAATGTTGAGATTTCCCGCGCTTGAGAATA |
| 5 | TCATCTCGTCAAACAATTTGGCCCAATTACAAGCGGCTTACGACGAAGCACTAGTGGCAA |

**Table S6.** T-test values and significance of each conjugate compared with the fluorescent reporter. The analysis was performed using SciPy: Open-Source Scientific Tools for Python. T-tests with P-values <0.05 were considered to be significant.

| | T-statistic | P-value | Significance |
|---|---|---|---|
| $^{Bio}DNA_{20}$ | -2.25 | 0.0872 | **Not significant** |
| $^{Bio}DNA_{40}$ | 0.87 | 0.8726 | **Not significant** |
| $^{Bio}DNA_{100}$ | 4.35 | 0.0122 | **Significant** |

**Table S7.** T-test values and significance of samples before, after and one week after lyophilization with DNA target vs. without. The analysis was performed using SciPy: Open-Source Scientific Tools for Python. T-tests with P-values <0.05 were considered to be significant.

| | T-statistic | P-value | Significance |
|---|---|---|---|
| Before | 10.16 | <0.0001 | **Significant** |
| After | 6.84 | 0.0001 | **Significant** |
| One week after | 4.86 | 0.0013 | **Significant** |

a)

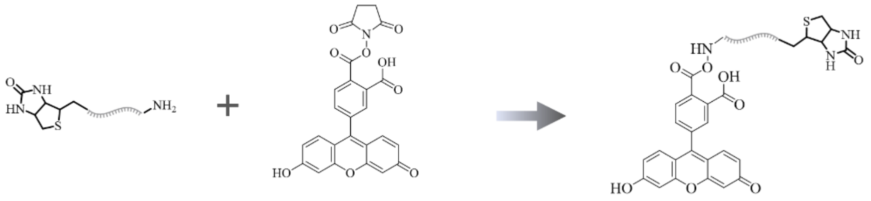


b)

| | 1x PBS | 5x PBS | 10x PBS |
|---|---|---|---|
| % FAM-NHS per DNA | 8.54 | 34.59 | 42.74 |

c)

| | 25 µl | 50 µl | 100 µl | 200 µl |
|---|---|---|---|---|
| % FAM-NHS per DNA | 38.26 | 51.54 | 52.26 | 51.455 |

d)

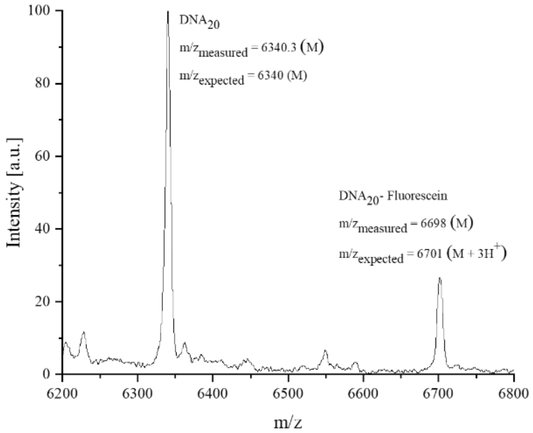


**Figure S1.** Aminylated DNA-NHS ester reaction optimization. a) Scheme of the reaction to obtain fluorescein conjugated to DNA. b) PBS concentration titration in 100 µl volume at pH 8.4 with 10 nmol of aminylated and 100 nmol FAM-NHS. c) Volume titration in 10x PBS at pH 8.4 with 10 nmol of aminylated and 100 nmol FAM-NHS. d) Mass spectrometry of the reaction in part b.

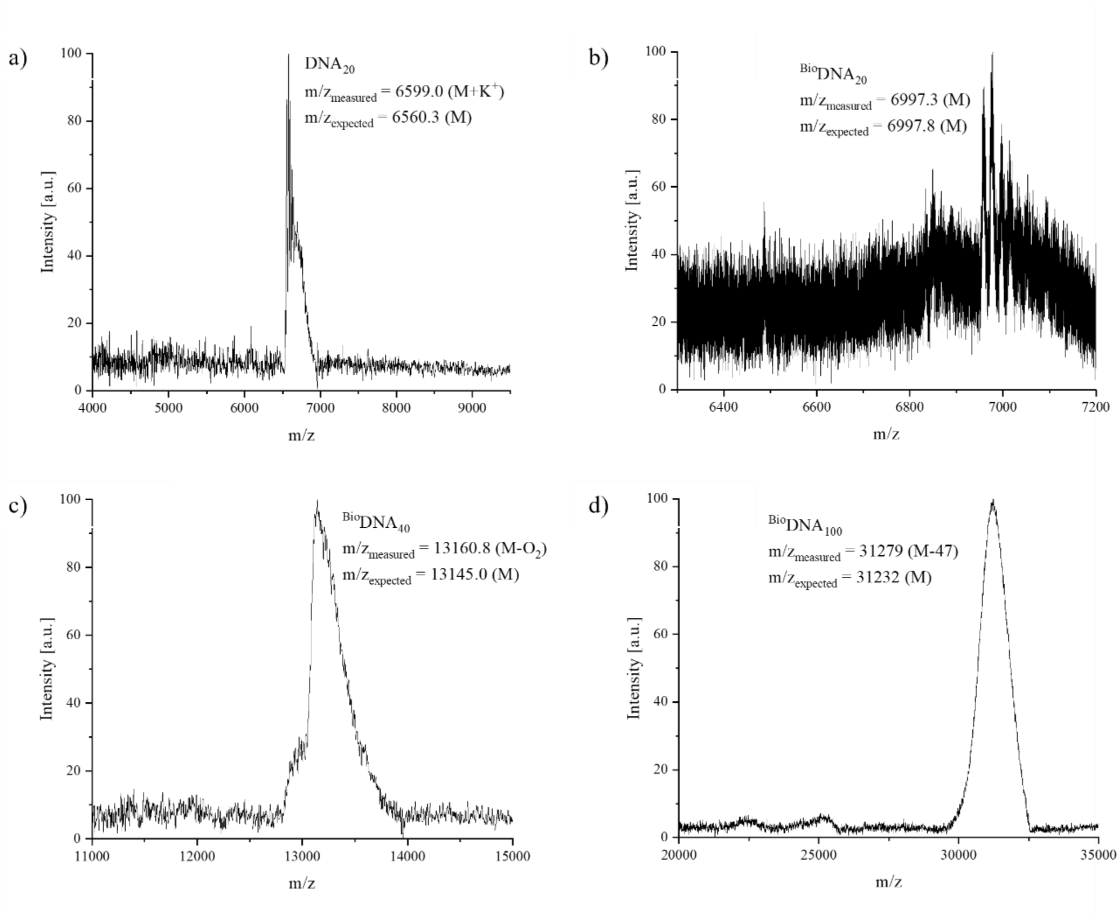


**Figure S2.** MALDI-TOFs of DNA-SMCC samples. a) 20 bp 5' aminylated DNA. b) 20 bp 5' aminylated and 3' biotinylated DNA. c) 40 bp 5' aminylated and 3' biotinylated DNA. d) 100 bp 5' aminylated and 3' biotinylated DNA.

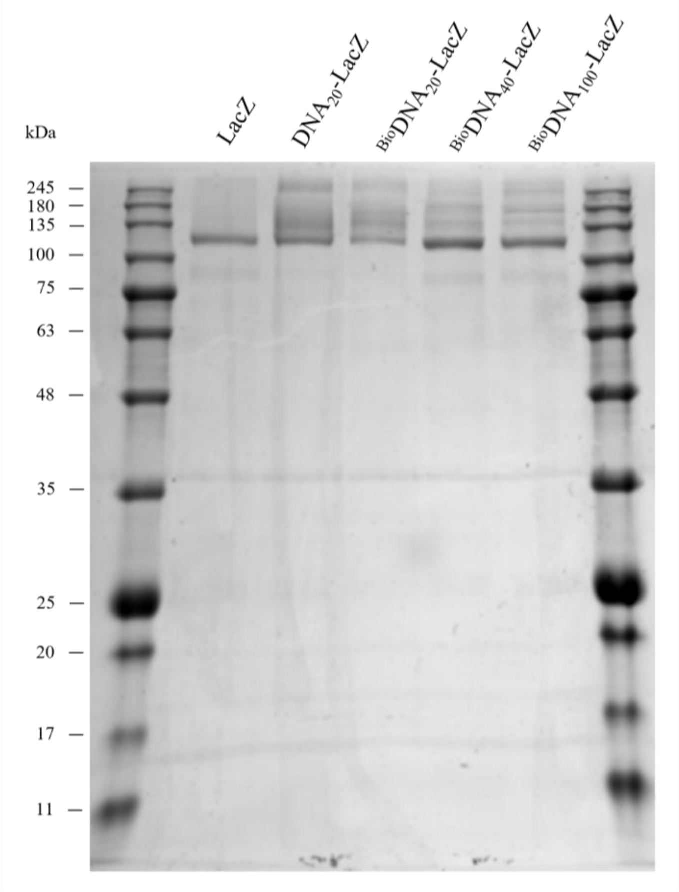


**Figure S3.** SDS-PAGE of LacZ at ~118 kDa vs. DNA-LacZ conjugates at the same and higher masses.

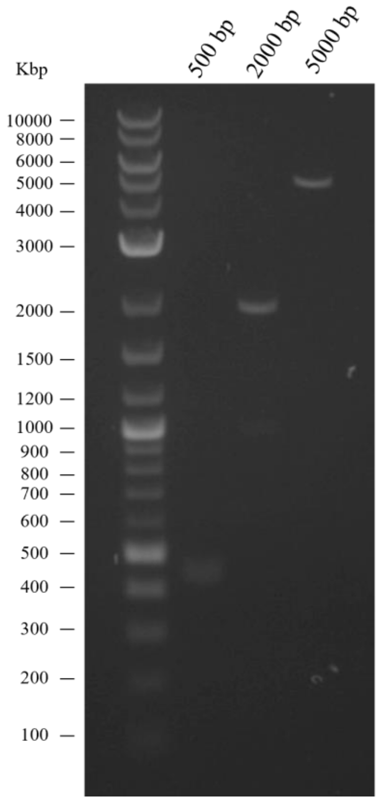


**Figure S4.** Gel electrophoresis of 500, 2000 and 5000 bp *S.* Typhi target amplicons.

For LacZ, it was clear that an additive was necessary since all activity was lost without one after freeze drying, while most additives tested allowed it to maintain its activity. It is important to note, that although LacZ remained active after freeze drying, all additives had some inhibitory effect. Cas12 could maintain some activity without an additive but was also inhibited by most of the compounds tested. The exceptions to this were D-Glucose, Sucrose and D-(+)-Trehalose dihydrate with trehalose being the highest performing one, and therefore, selected for the full system.

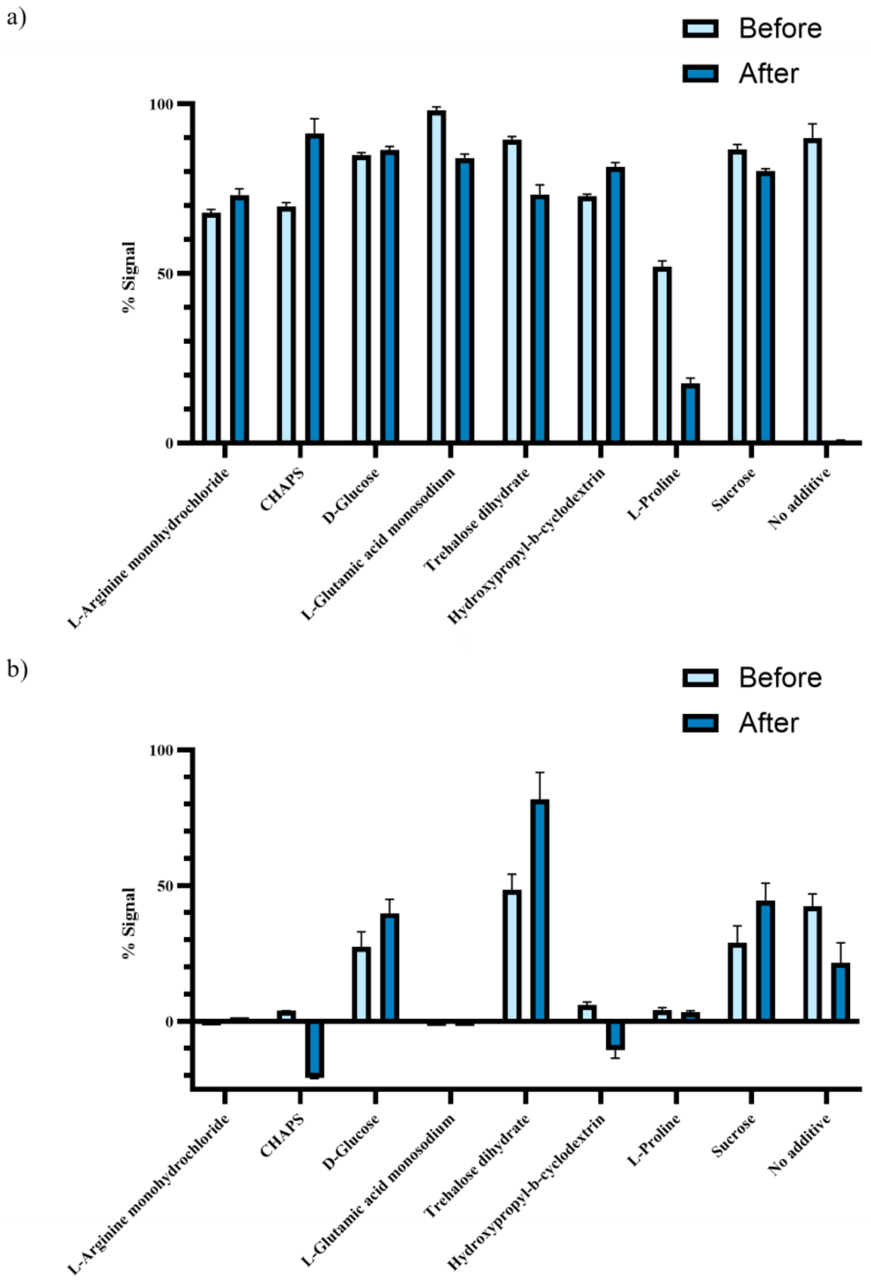


**Figure S5.** Additive effect on enzyme activity before and after lyophilization. a) 10% w/v additives mixed with 0.8 µg LacZ. Enzyme rehydrated to the original volume with water and mixed with X-Gal at a final solution of 2.5 mg/ml. Absorbance measured after 20 minutes of incubation at 37 °C, subtracted by the signal at time zero and divided by the highest absorbance value across all conditions. b) 10% w/v additives mixed with 75 nM LbaCas12a, 95 nM gRNA and 1x CutsmartT. Components rehydrated to the original volume with water and mixed with 10 nM 60 bp synthetic DNA target and 0.5 µM FQ reporter. Fluorescence measured after 20 minutes of incubation at 37 °C, subtracted by the signal at time zero and divided by the highest fluorescence value across all conditions.
